# Benchmarking ensemble docking methods as a scientific outreach project

**DOI:** 10.1101/2020.10.02.324343

**Authors:** Jessie L. Gan, Dhruv Kumar, Cynthia Chen, Bryn C. Taylor, Benjamin R. Jagger, Rommie E. Amaro, Christopher T. Lee

## Abstract

The discovery of new drugs is a time consuming and expensive process. Methods such as virtual screening, which can filter out ineffective compounds from drug libraries prior to expensive experimental study, have become popular research topics. As the computational drug discovery community has grown, in order to benchmark the various advances in methodology, organizations such as the Drug Design Data Resource have begun hosting blinded grand challenges seeking to identify the best methods for ligand pose-prediction, ligand affinity ranking, and free energy calculations. Such open challenges offer a unique opportunity for researchers to partner with junior students (e.g., high school and undergraduate) to validate basic yet fundamental hypotheses considered to be uninteresting to domain experts. Here, we, a group of high school-aged students and their mentors, present the results of our participation in Grand Challenge 4 where we predicted ligand affinity rankings for the Cathepsin S protease, an important protein target for autoimmune diseases. To investigate the effect of incorporating receptor dynamics on ligand affinity rankings, we employed the Relaxed Complex Scheme, a molecular docking method paired with molecular dynamics-generated receptor conformations. We found that CatS is a difficult target for molecular docking and we explore some advanced methods such as distance-restrained docking to try to improve the correlation with experiments. This project has exemplified the capabilities of high school students when supported with a rigorous curriculum, and demonstrates the value of community-driven competitions for beginners in computational drug discovery.

## 1 INTRODUCTION

Drug discovery efforts often require the screening of many compounds to determine their efficacy. Owing to the high cost of experimental screening and advances in computer models, the use of inexpensive computational screening methods to enrich compounds in large datasets have been used in drug discovery pipelines for several decades (1). Early on in the screening process, when an initial compound library may contain only a few ‘active’ among many orders of magnitude more inactive compounds, computer-aided drug discovery (CADD) methods, such as virtual screening, can be used to filter out unlikely candidates, reducing experimental costs, and accelerating the initial discovery phase (2–4).

Due to the diversity and breadth of the CADD research community, many methods have been developed. Cross-comparison and benchmarking between the different approaches is necessary for identifying the limitations of the docking method and areas for improvement. The Drug Design Data Resource (D3R) hosts blinded community prediction challenges to evaluate these software and techniques and compare their effectiveness on benchmark systems, such as the HSP90 chaperone protein, the Farnesoid X nuclear receptor, and the Cathepsin S protease (CatS) (5–7). In 2018, D3R hosted Grand Challenge 4 (GC4), which had components of pose prediction, free energy prediction, and ligand affinity rank ordering (8).

We participated in Subchallenge 2, a ligand affinity ranking challenge for the Cathepsin S protease with a set of 459 ligands provided by Janssen Pharmaceuticals (8). CatS is a cysteine protease involved in the presentation of antigens by the MHC class II molecules within CD4^+^ T cells (9). This makes it a promising target in autoimmune disease and allergy treatment, where inhibition of the immune response is critical for effective therapy (10–12).

We used molecular docking, a popular method of virtual screening, in a strategy known as the Relaxed Complex Scheme to account for protein flexibility (13). Molecular docking applies a conformational search algorithm paired with an inexpensive, and often empirical, scoring function to find favorable lead compounds (14, 15). By forgoing rigorous dynamics and detailed potential energy functions, such as those used in free energy calculations, docking approaches are designed to yield results quickly albeit with lower accuracy (16). The speed of molecular docking codes enables the screening of hundreds of thousands to millions of compounds (17). A risk of docking is the increased likelihood of false negatives. To this end, much work has been done by the community to develop improved algorithms which improve docking accuracy with minimal impact on speed (3).

In early docking studies, proteins and ligands were represented as static structures (16, 18, 19). To incorporate ligand flexibility, multiple ligand positions can be sampled through rotational torsions, i.e. conformer generation (20). However, Molecular Dynamics (MD) simulations have revealed that thermal protein fluctuations in solute-based environments can give rise to varying conformational states, resulting in different binding sites (21). Accounting for receptor binding site flexibility in molecular docking is a significant challenge. One solution is to perform ensemble docking. This involves docking a ligand compound library to a number of distinct, rigid receptor conformations to identify the receptor conformation that is best suited for that particular ligand (i.e. best docking score) (20, 22–24).

Here, we perform MD simulations of the receptor protein, CatS, to obtain unique conformational states and introduce structural variation in the binding site. MD simulations allow the exploration of multiple conformations of the protein while in a solute-based, native environment (25, 26). This concept of selecting naturally-occurring conformations through MD for ensemble docking is known as the Relaxed Complex Scheme (26–31). MD-generated ensembles of flexible binding sites have been used successfully in a number of studies to identify lead compounds (13, 32–35).

Incorporating more receptor conformations increases computational cost, as a complete docking protocol must be performed for each conformation. To address this, the trajectory can be clustered to extract unique, representative conformations (36, 37). This methodology is still susceptible to the conformational sampling problem of MD, due to the large discrepancy between the accessible timescales of MD simulation (microseconds) and the slow, native dynamics of proteins (milliseconds and longer) (38, 39). Although a trajectory may not statistically converge to encompass all possible conformations, studies have shown that clustering MD trajectories can reveal previously unknown druggable pockets (32).

Many studies have successfully used clustering methods in ensemble docking to extract relevant conformations, such as those based on RMSD (26, 34), QR factorization (13, 33), and active pocket volume (26). However, choosing the most appropriate clustering method for a system is still challenging and often dependent on human intuition.

Although ensemble docking has been successfully used to identify lead compounds, clustering methods in ensemble docking have not been extensively studied (40). We explored three clustering methodologies in this study to investigate if they could (i) provide an accurate ligand ranking and (ii) give insights into CatS ligand binding mechanisms. The three clustering methods we used are: 1) Time-lagged Independent Components Analysis and K-means clustering (TICA) (41, 42), 2) Principal Component Analysis and K-means clustering (PCA) (41, 42), and 3) Gromos RMSD clustering (Gromos) (43). TICA identifies the slowest motions of the simulation and projects the input features into a slow subspace where distinct clusters are kinetically separated (44). PCA, on the other hand, finds features with the largest variance (45). Lastly, Gromos is a RMSD-based clustering method that counts the neighbors in a cluster based on a pre-set cutoff value and defines trajectory clusters by structural variation (43).

In this work, we apply the Relaxed Complex Scheme with these clustering methods and compare the ensemble docking results (30). We test the accuracy of two state-of-the-art docking softwares: Open Eye FRED (46) and Schrodinger Glide (47). We found that CatS is a difficult target for molecular docking and we explore some advanced methods such as distance-restrained docking to try to improve the correlation with experiments.

## 2 PEDAGOGICAL SIGNIFICANCE

This manuscript presents the work of high school students who have performed this work after completing BioChem-CoRe, a 7 week crash course on computational chemistry (http://biochemcore.ucsd.edu/). These results helps to illustrate the benefits and possibilities of teaching science as we do science (48, 49). By participating in structured challenges with real-world significance, students gain motivation, confidence, and both technical and soft skills. Moreover, the exposure to the rigors of the scientific approach and the methods employed in the field of study aids them with their future career decisions. On the other hand, community-driven competitions and resources such as D3R’s Grand Challenge 4 can also benefit from student participation. Rarely do these programs receive submission which test the basic hypothesis. For example, is domain expertise required for the application of the methods of interest? Given the current state of tutorials or instructions available to the public, can students with limited domain experience use these resources to produce results without major technical difficulties? We posit that student participation can not only yield important benchmarking data but also serve to improve the documentation of our tools and methods.

## 3 MATERIALS AND METHODS

All scripts used in this work can be found online at https://github.com/ctlee/bccgc4. Full workflow of methods is shown in Fig. 1.

**Figure 1:**
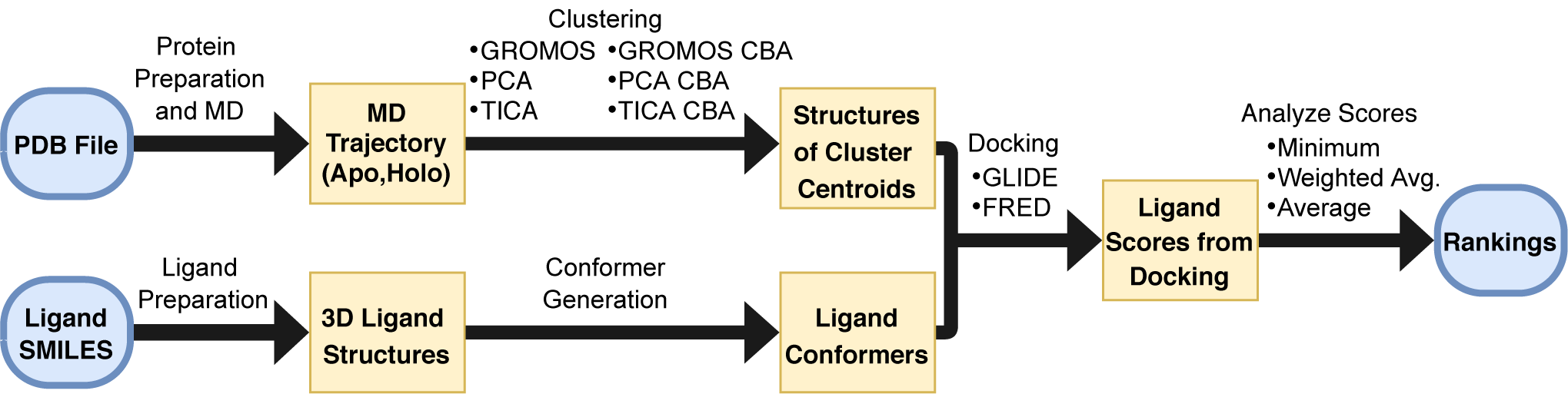
Workflow of ensemble docking approach. PDB file was selected and simulated in Molecular Dynamics. Molecular Dynamics trajectory was clustered by six various methods and cluster centroids were extracted as representative structures. Ligand SMILES were prepared as 3D structures and various conformers were generated. Molecular docking of ligands to cluster centroids was performed with FRED and Glide docking. Pose scores were used to generate rank orderings and Kendall’s *τ* values when compared to the experimental rank ordering.

### 3.1 Molecular Dynamics

A crystal structure of CatS (PDBID: 5QC4 (11)) was obtained from the RCSB PDB database (50). The structure was chosen due to its resolution of 2 Å and similarity of the cocrystallized ligand to those in the D3R dataset. This cocrystal was part of D3R’s prior Grand Challenge 3 (GC3), subchallenge 1, and the ligands in the CatS subchallenge of Grand Challenge 4 all contain the tetrahydropyrido-pyrazole core that 22 of the 24 ligands had in the previous challenge (7).

Models with (holo) and without (apo) the cocrystallized ligand were prepared for MD simulations. For both apo and holo models the same steps were performed with a few deviations noted below. Chain A of the structure was prepared in Schrodinger Maestro 2019 (51) with the Protein Preparation Wizard. For the holo simulation, the cocrystallized ligand was retained. For the apo simulation the ligand was removed (52). Force field parameters for the ligand were derived from GAFF (53) with partial charges fit using the restrained electrostatic potential method (RESP) (54) from potentials computed using the AM1-BCC semi-empirical quantum mechanical method (55, 56). For both systems, the protein termini were capped with an acetyl (ACE) and N-methyl amide (NME) capping groups. PROPKA (57, 58) was used to assign residue protonation states in a solvent of pH 5.0, to mimic experimental conditions of CatS binding assays (9). Crystal waters with more than 2 hydrogen bonds to non-waters were retained.

Using a combination of pdb4amber and tleap from the AMBER 18 software suite, we parameterized the systems with the AMBER FF14SB forcefield, and solvated the systems with TIP4P-Ew up to a 15 Å buffer distance (59). We added ions according to the SLTCAP tool by Schmit et al. (60) at 100 mM salt concentration, again to mimic experimental conditions (9).

All-atom, explicit-solvent MD simulation was performed for the both systems using AMBER18 in four stages: minimization, heating, equilibration, and production (59). The systems were gradually minimized in four steps: (i) minimization of only protons, restraining the protein and solvent, (ii) minimization of the solvent, restraining the protein, (iii) minimization of the protein sidechains, restraining the protein backbone, and (iv) minimization of all atoms. Restrained heating was performed in two steps: first, in the NVT ensemble the temperature was increased from 0 to 100 K over 50 ps using a Langevin thermostat, and second in the NPT ensemble the temperature was increased from 100 to 300 K over 200 ps using a Langevin thermostat while pressure was maintained at 1 bar using a Berendesen barostat. Equilibration was also performed in two stages, first with a restrained backbone, and second without restraints. For both equilibration stages the temperature was maintained at 300 K using a Langevin thermostat. For the restrained equilibration stage, 500 ps were run with a Berendsen barostat to equilibrate pressure to 1 bar. In the unrestrained equilibration step 1000 ps were run using a Monte Carlo barostat at 1 bar.

Production simulations were run in the NPT ensemble with the same conditions as the unrestrained equilibration step. Five independent simulations of each condition, apo and holo, of length 2 *µ*s were run, totaling 20 *µ*s. Hydrogen Mass Repartitioning (HMR) was performed with PARMED (59, 61) for all systems permitting a 4 fs timestep. All simulations were run with SHAKE restraints (62) and a non-bonded cutoff of 10 Å.

### 3.2 Clustering

The MD trajectory was clustered using three different clustering methods: 1) TICA and k-means (41, 42, 63, 64) on the protein backbone atom position coordinates, 2) PCA and k-means on the protein backbone atom position coordinates, and 3) Gromos (43) on the C-alpha atom position coordinates. To identify a good set of initial input features, we compared the mean 10-fold cross-validated Variational Approach for Markov Processes (VAMP2) scores for three selections: i) protein backbone atom positions, ii) protein backbone torsions, and iii) the positions of a binding atoms selection (65). We decided to use the positions of protein backbone atoms because it had the largest VAMP2 score, indicating greater kinetic variance. The binding atoms were defined by taking all receptor atoms within 2 Å of the initial docked poses of a ligand from the D3R data-set. PCA was clustered on the same subset of backbone atom positions (41), and Gromos was clustered on the C-alpha positions due to memory limitations. After the challenge, the clustering was reevaluated and a second discretization using the binding atoms selection, referred to as Clustered by Binding Atoms (CBA), was generated. We used similar ideas as the approach taken in Ref. (66), focusing on the binding site’s structural fluctuations rather than the entire structure. All six clustering methods (TICA, PCA, and Gromos for backbone or C-alpha atoms and CBA) were also performed on the holo MD trajectories. The cluster centroids of the apo MD were compared by pairwise RMSD, utilizing MDTraj and NumPy together to calculate RMSDs and order them into a matrix which was visualized in matplotlib (67–69). They were also compared in terms of Root-Mean-Squared-Fluctuation (RMSF) to investigate the particular structural variability, computed in MDTraj and visualized in PyMOL (68, 70).

#### 3.2.1 Time-lagged Independent Components Analysis and K-means (TICA)

TICA clustering was employed to capture the slow motions within the trajectory. TICA was performed with a lag time of 4 ps and a variance cutoff of 0.95 on the protein backbone atom coordinates (41). The trajectory was projected into the TIC basis and subsequently, the k-means algorithm was used to cluster the trajectory into 10 distinct clusters. The 10 configurations from the trajectory, in real space, closest in TIC space to the cluster centroids were used for docking (42).

#### 3.2.2 Principal Components Analysis and K-means (PCA)

PCA with a variance cutoff of 0.95 was performed on the protein backbone atom coordinates to capture large motions within the trajectory (41). The trajectory was projected into the PC basis and subsequently, the k-means algorithm was used to cluster the trajectory into 10 clusters. The 10 configurations from the trajectory, in real space, closest in PC space to the cluster centroids were used for docking (42).

#### 3.2.3 Gromacs RMSD-Based Clustering (Gromos)

Gromos clustering was performed on the alpha carbons in the protein to identify structurally diverse conformations according to RMSD (43). The trajectories input to Gromos were subsampled to yield frames every 0.4 ps. This was due to computational intractability at more frequent frame rates. The clustering RMSD cutoff was chosen to satisfy the following criteria: (i) the first cluster had less than 70% of the frames, (ii) the first 10 clusters contained at least 80% of the frames, and (iii) each of the first 10 clusters had at least 20 frames. A cutoff of 0.08 Å was used when clustering with alpha carbons while a cutoff of 0.15 Å was used when compared to CBA for the apo trajectory. A cutoff of 0.07 Å was used when clustering with alpha carbons while a cutoff of 0.135 Å was used when compared to CBA for the holo trajectory.

### 3.3 Docking

OpenEye Scientifics Fast Exhaustive Docking (FRED) was used to dock the 459 ligands in the GC4 CatS challenge to the centroids of the clustered MD trajectory and the original crystal structure (46). After the challenge, Schrodingers Glide was also used in an attempt to improve rank ordering and pose prediction (47, 71). In addition, many iterations of Glide docking were run with modifications to further improve the results. The pose results were visualized in Schrodinger’s Maestro (51) and labeled in Inkscape (72). The pose results were analyzed for accuracy through the RMSD of the common core to the original cocrystal ligand core, calculated using Schrodinger’s Python API and visualized in matplotlib (69).

#### 3.3.1 FRED

OpenEye’s OMEGA was used to convert ligand SMILES to 3D conformers, with the maximum number of conformers per ligand set to 800 (73). The conformers were then docked to the crystal structure and the 10 cluster centroids from each clustering method using OEDockings FRED default settings (Chemgauss4 scoring function with standard search resolution) (46, 74). The receptor area was defined by a box around the protein, determined by the minimum and maximum distance coordinates of the entire protein. For each receptor ensemble, the minimum score of every ligand was used in determining the rank ordering, as in previous studies (13, 32).

#### 3.3.2 Glide

Schrodingers Ligprep was used to convert ligand SMILES using standard settings into Maestro structures for Schrodingers Glide docking (75). Glides cross-docking script, xglide.py, was used to perform ensemble docking for each clustering method. The cross-docking script generated receptor grid files for each centroid structure using a 32 Å box centered on the center of mass of the crystal structure’s ligand (BC7 (11)) to define the docking region. Each centroid was then docked to using Glides Standard Precision (SP) docking methodology, which has its own ligand conformer generation steps, and scored with the subsequent Standard Precision GlideScore scoring function (47, 71). For each ensemble docking approach, the best score of each ligand across the ensemble of conformations (**N**) was used to determine its rank,

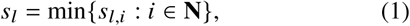

where *s*_*l*_ and *s*_*l,i*_ are the best overall score and best score for receptor conformation *i* for ligand *l* respectively.

To further investigate the ligand binding we also 1) applied a restraint on the tetrahydropyrido-pyrazole common core structure, restricted to lie within 3.5 Å of the cocrystal ligand’s common core, 2) changed the precision of the docking and scoring function from Glide SP to Glide Extra Precision (XP) (76), and 3) clustering and docking to centroids from a holo MD trajectory.

### 3.4 Scoring Schemes for Ligand Scores

Aside from changing the docking methodology, we also tried two other scoring schemes such as taking the average and the weighted average (Eq. (2)) of the OpenEye’s FRED and Glide SP scores.

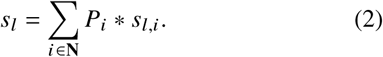

where *P*_*i*_ is the probability of observing conformation *i, s*_*l,i*_ is the best docked score for that conformation, and **N** is the set of conformations in the ensemble. Note that the probabilities, *P*_*i*_, are normalized such that Σ_*i*∈**N**_ *P*_*i*_ = 1. The *P*_*i*_ for a given conformation *i* is calculated as *f*_*i*_ *f*_*T*_, where *f*_*i*_ is the number of frames in the same cluster as *i*, and *f*_*T*_ is the total number of frames in the trajectory. These scoring schemes have been used in other studies due to the reasoning that the average (33, 77) or weighted average (34, 35) score better accounts for the variability of the ensemble, and in the case of the weighted average, represents the likelihood of the ligand encountering each representative conformation in a natural environment.

### 3.5 Kendall’s Taus

Ligand rankings were created by sorting the ligands based on their score. Kendall’s Tau values were calculated by comparing the predicted rank ordering to the experimental rank ordering using the Kendall’s Tau function in SciPy (78).

## 4 RESULTS AND DISCUSSION

In lieu of running expensive free energy calculations which account for both ligand and receptor flexibility, the Relaxed Complex Scheme attempts to reduce computational cost while capturing the flexibility of a protein by docking to multiple protein conformations selected from a MD simulation. These representative conformations are often chosen by combining a method of dimensionality reduction followed by the application of a clustering algorithm. Although this approach is conceptually simple, the choice of clusters has many pitfalls. For example, even if a set of clusters spans the conformational diversity of the MD trajectory, the ensemble will not necessarily produce the most accurate ligand rank ordering (40). Some receptors may have natural conformations which are not ideal for ligand binding, and these may result in false positives (25). In addition, the active conformation for ligand binding could be transient, and would have a lower probability of being represented in the ensemble.

To test the effect of clustering approach on the resulting conformations, we test several different clustering methods. TICA captures slow protein movements (variance in time), while PCA focuses on large structural variance, and Gromos captures structural variations as measured by RMSD. We plot the structural variation across clusters from the different algorithms in Fig. 2B,C, where CBA refers to clustered by binding atoms, defined in Fig. 2A. We find that centroids from different clustering methods vary in different structural domains, Fig. 2C. The structural fluctuation around the binding site (facing the reader in Fig. 2C) are most likely to affect ligand binding. By restricting the set features input to the clustering workflow to the binding atoms, we find that the CBA methods capture increased variability in the binding site.

**Figure 2:**
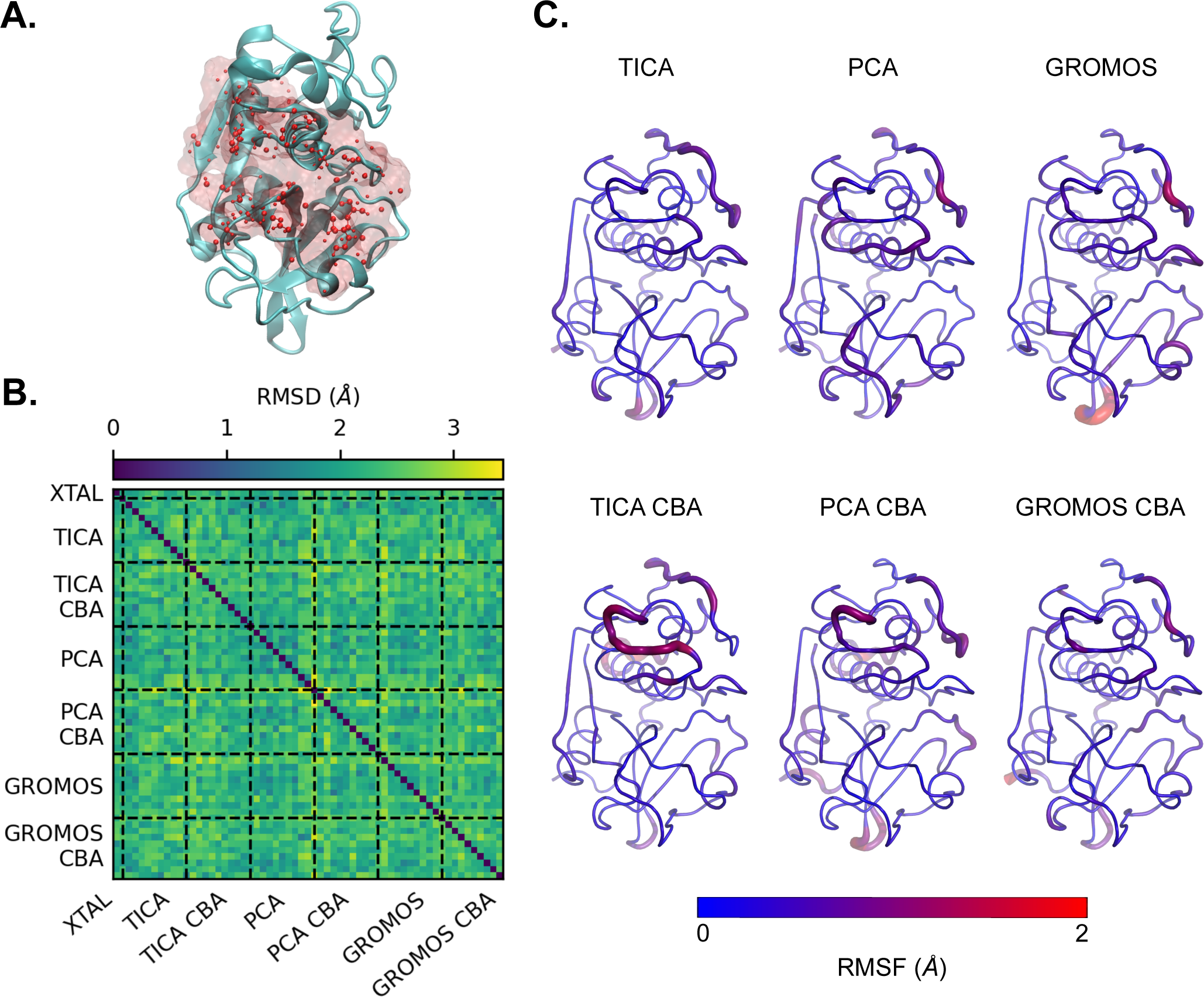
Apo Molecular Dynamics (MD) Clustering Results. A) Binding Atoms definition for Clustered by Binding Atoms (CBA) centroids, defined by taking all atoms within 2 Å of docked poses of a ligand from the D3R dataset (CatS_2) from both Glide and FRED SP apo blind docking. The crystal structure protein is depicted in NewCartoon and colored teal, while the binding atoms are both represented by red spheres and a transparent red surface representation, visualized in Visual Molecular Dynamics (VMD) (79, 80). B) The pairwise Root-Means-Squared-Deviations (RMSDs) of the binding atoms of the crystal structure and all 10 centroid structures from each clustering method are depicted in a heatmap. The centroids obtained from clustering have a range of RMSDs and therefore have structural variability. C) MD clustering extracts various centroid structures, and different clustering methods yield different conformations. The RMSF of the 10 centroids extracted from each clustering method, shown as the relative thickness and color, was calculated with MDTraj (68) and visualized using PyMOL (70). The orientation of the protein for parts A and C are the same.

### 4.1 Initial Docking

Our FRED docking results performed worse than random rank ordering (Fig. 3A). To investigate the influence of docking algorithm, both scoring and conformational searching, we also performed docking with Schrodinger’s Glide (81). The rank order correlation of the predictions from Glide docking were better than random rank ordering (Fig. 3B).

**Figure 3:**
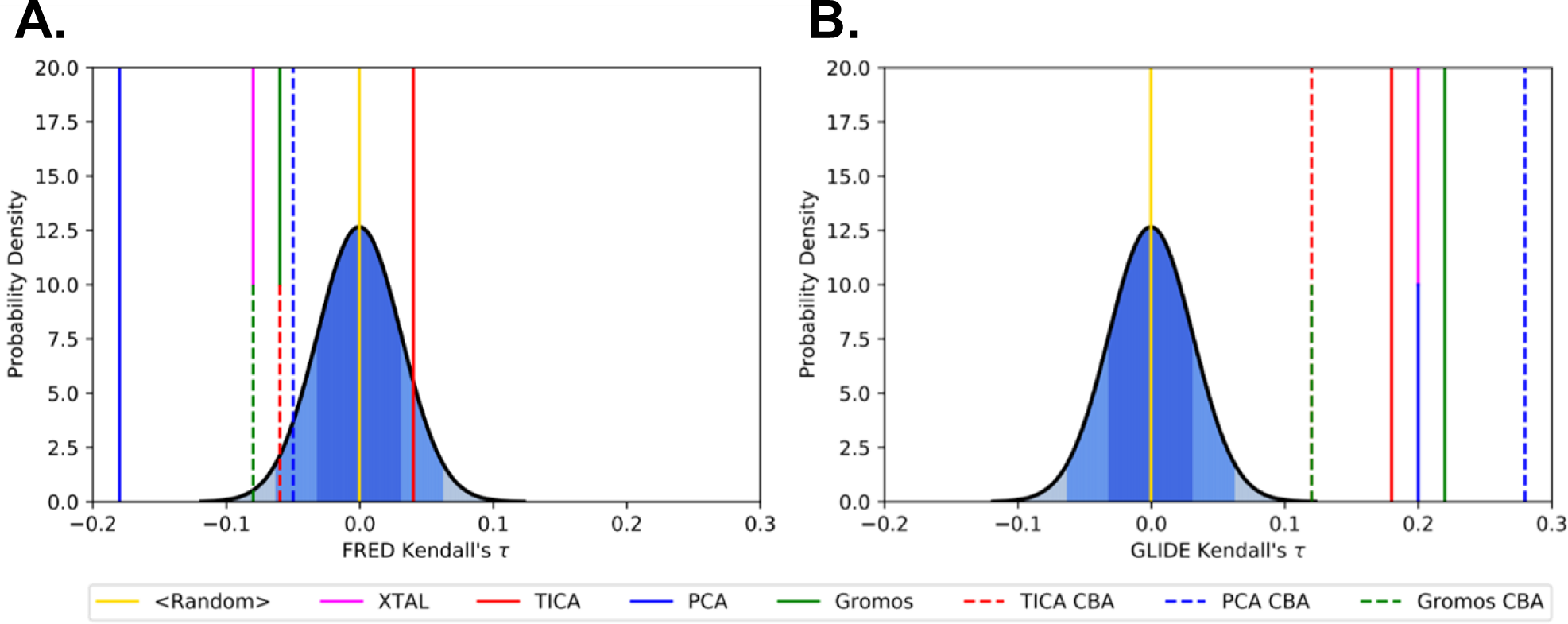
The FRED rank ordering results were unsuccessful in producing a higher Kendall’s *τ* value than random ordering, while the Glide results were able to predict better than random. A) Kendall’s *τ* values for ligand rankings based on minimum scores from OpenEye’s FRED blind docking to apo MD centroids, compared to a random rank ordering distribution. B) Kendall’s *τ* values for ligand rankings based on minimum scores from Schrodinger’s Glide docking to apo MD centroids, compared to a random rank ordering distribution. In both A) and B) a probability distribution function is graphed from the Kendall’s *τ* values of 10,000 random ligand rank orderings. The distribution has *µ* = 0 and *σ* = 0.031.

There are some inherent differences between OpenEye’s FRED and Schrodinger’s Glide conformational search algorithms. FRED’s docking algorithm emphasizes shape complementarity between the ligand and protein through an exhaustive pose search that samples multiple ligand positions. It accounts for ligand rotations and scores multiple poses before selecting one top scoring pose per ligand (46). On the other hand, Glide’s docking algorithm begins with receptor grid generation and focuses on ligand binding energy, including a ligand minimization with a standard molecular mechanics energy function, the OPLS-AA force field, and a distance-dependent dielectric model. In addition, the final poses are refined with a Monte-Carlo procedure to find torsional minima (47). Both methods consider ligand conformers, either generated separately (through OpenEye OMEGA) or as part of the docking workflow (Glide).

The scoring functions also differ between the two software. For FRED, the ChemGauss 4 scoring function, which uses Gaussian-smoothed step-function based interaction potentials, is used to optimize top poses from the filtering steps (46). Meanwhile, Glide’s GlideScore uses more complex and varied weight functions for the various potential terms (47). The more complex approach to fit empirical scoring functions used by Glide may have improved the pose prediction similarity to cocrystal poses and the rank ordering accuracy.

### 4.2 Scoring Scheme Results

Next, we investigated if the approach to compute a single score from an ensemble of scores can improve the accuracy of our predictions. There are several ways to obtain a single score from an ensemble of values. The first is to take the minimum score of the ensemble. This assumes that the other configurations do not contribute to the ligand binding energy. Relaxing this assumption, it is possible to consider the contributions of other receptor configurations by using an average or weighted average of the ensemble values. The choice of weights may be assigned by the probability of observing each conformation among other strategies. Limitations from the limited sampling of MD may lead to unintended biases in the ensemble weights.

In our results, we saw minor fluctuations in Kendall’s *τ*s across different scoring schemes (Table 1). While some conditions saw improvements to Kendall’s *τ* when using the weighted average versus the minimum score, no consistent rationale for these improvements were found. It is therefore unclear from this system and study whether or not incorporating receptor flexibility can improve predictions of rank ordered correlation. We hypothesize that the challenges of docking to CatS which has a large solvent-exposed binding pocket may outweigh the benefits of incorporating receptor flexibility which has been reported in other works (26, 30).

**Table 1:** The Kendall’s *τ*s for the FRED and initial Glide docking show slight fluctuations in different scoring schemes, but do not show any immense improvement. Here we show the Kendall’s *τ* from rank orderings produced through various docking functions, clustering methods, and scoring schemes. Docking Functions are labeled accordingly: SP: Glide Standard Precision Docking, XP: Glide Extra Precision Docking; A: apo structure, H: holo structure; B: blind docking, R: restrained docking. We experimented with these scoring schemes to test if a particular method of discerning scores for each ensemble would better represent the protein binding mechanisms and improve rank ordering. The various scoring schemes were the Minimum, Weighted Average (W. Avg.), and Average (Avg.).

| Kendall's $\tau$ For All Ligand Rankings | | | | | | | | |
| --- | --- | --- | --- | --- | --- | --- | --- | --- |
| Docking Function | Scoring Method | Clustering Methods |  |  |  |  |  |  |
|  |  | XTAL | TICA | PCA | GROMOS | TICA CBA | PCA CBA | GROMOS CBA |
| FRED AB | Minimum | -0.08 | 0.04 | -0.18 | -0.06 | -0.06 | -0.05 | -0.08 |
|  | W. Avg. | - | -0.11 | -0.20 | -0.04 | -0.11 | -0.09 | -0.06 |
|  | Avg. | - | -0.13 | -0.09 | -0.10 | -0.09 | -0.21 | -0.10 |
| Glide SP-AB | Minimum | 0.20 | 0.18 | 0.18 | 0.22 | 0.12 | 0.28 | 0.12 |
|  | W. Avg. | - | 0.21 | 0.20 | 0.18 | 0.17 | 0.24 | 0.21 |
|  | Avg. | - | 0.20 | 0.21 | 0.25 | 0.20 | 0.24 | 0.23 |
| Glide SP-AR | Minimum | 0.13 | 0.14 | 0.13 | 0.13 | 0.11 | 0.11 | 0.09 |
|  | W. Avg. | - | 0.08 | 0.07 | 0.10 | 0.09 | 0.05 | 0.07 |
|  | Avg. | - | 0.12 | 0.07 | 0.09 | 0.07 | 0.06 | 0.09 |
| Glide XP-AB | Minimum | 0.20 | 0.11 | 0.11 | 0.12 | 0.11 | 0.24 | 0.14 |
|  | W. Avg. | - | 0.10 | 0.08 | 0.08 | 0.11 | 0.17 | 0.15 |
|  | Avg. | - | 0.11 | 0.08 | 0.07 | 0.12 | 0.19 | 0.17 |
| Glide XP-AR | Minimum | 0.13 | 0.14 | 0.10 | 0.12 | 0.11 | 0.09 | 0.13 |
|  | W. Avg. | - | 0.10 | 0.07 | 0.04 | 0.07 | 0.03 | 0.11 |
|  | Avg. | - | 0.11 | 0.07 | 0.06 | 0.04 | 0.04 | 0.08 |
| Glide SP-HB | Minimum | 0.09 | 0.17 | 0.14 | 0.18 | 0.23 | 0.23 | 0.18 |
|  | W. Avg. | - | 0.20 | -0.01 | 0.17 | 0.24 | 0.14 | 0.20 |
|  | Avg. | - | 0.22 | -0.01 | 0.21 | 0.23 | 0.18 | 0.21 |
| Glide SP-HR | Minimum | 0.12 | 0.18 | 0.13 | 0.11 | 0.16 | 0.17 | 0.15 |
|  | W. Avg. | - | 0.13 | 0.09 | 0.08 | 0.15 | 0.11 | 0.13 |
|  | Avg. | - | 0.15 | 0.11 | 0.12 | 0.14 | 0.13 | 0.14 |

To further understand the shortcomings in our approach, we conducted multiple revisions to both the trajectory clustering and the docking methodology.

### 4.3 Pose Analysis and Glide Docking Revisions

We found that the ligands in the CatS dataset had a common tetrahydropyrido-pyrazole core to other ligands with published cocrystal structures from a prior D3R Grand Challenge (GC3) (Fig. S1) (7, 50). The poses from FRED docking were varied and often located opposite from the binding location of similar cocrystallized ligands (Fig. 4A). Other cocrystals contain ligands bound to this alternative site, although these ligands are dissimilar to the ones in our dataset (ligands 29 to 48 in Fig. S1, Table. S1) (82).

**Figure 4:**
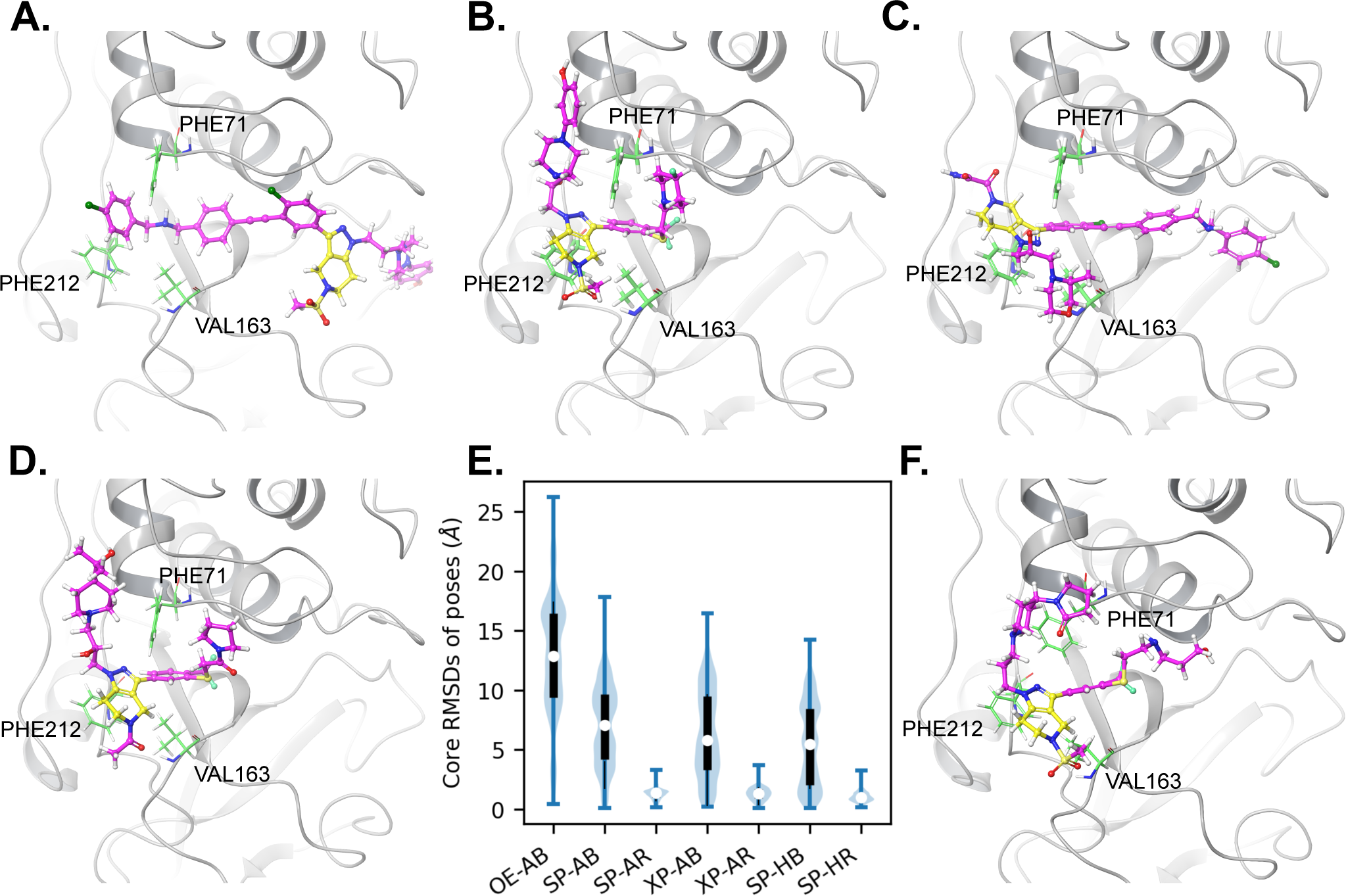
Docking pose analysis shows that a distance-restraint improves pose accuracy. A) Ligand CatS 259, example of an inaccurate FRED pose, with the core in a different location than the crystal structure of a similar ligand. B) Ligand CatS 118 of the SP apo blind crystal docking: ideal pose most similar to the cocrystal structure. C) Ligand CatS 363 of the SP apo blind crystal docking: some docked ligands show a flipped core binding mode that is less common but can be found in some available cocrystals. (7). D) cocrystal pose (PDBID: 5QC4 (11)) Ligand carbons are pink; ligand common core carbons are yellow; key binding residues PHE71, VAL163, and PHE212 are green. E) The RMSDs of the ligand core for each pose in each Glide docking method show that blind poses were concentrated farther from the cocrystal position compared to the ligand-core-restrained docking. In addition, the FRED average core RMSD is larger than that of all blinded Glide RMSDs. Each violin is composed of all minimum poses for each clustering method which contributed to the final rank ordering and the crystal structure poses, totaling *n* = 3213 per violin. Method Acronyms: OE: OpenEye FRED docking, SP: Glide Standard Precision Docking, XP: Glide Extra Precision Docking; A: apo structure, H: holo structure; B: blind docking, R: restrained docking. The median is represented in white, the interquartile range is shown in black, and the minimum and maximum values are shown as whiskers. F) Ligand CatS 23 of the SP apo restrained PCA docking: When the ligand is restrained, it can be unnaturally docked in receptors that are dissimilar to the cocrystal, such as here where the PHE71 is in a different configuration.

Glide docking produced some poses similar to the cocrystal pose (Fig. 4B) although it also produced more unexpected poses. We also observed cocrystals with ligands binding in the less common “flipped core” configuration, shown in Fig. 4C, reported in GC3 (7).

To test the hypothesis whether improved pose similarity to cocrystal structures can improve docking accuracy, we applied a distance-restraint to the common core of the ligands using the core position of published cocrystals with similar ligands as a reference point (Fig. 4D). Other work has found that approaches which use information from cocrystals such as template docking or restraints can improve pose accuracy (5–7, 83, 84). The restraint employed eliminated poses which deviate significantly from the cocrystal pose while permitting the flipped configuration. As shown in Fig. 4E, the RMSDs of the tetrahydropyrido-pyrazole core in SP apo docking were reduced (from a median of 7.39 Å to 1.40 Å) by adding the restraint, however, this did not improve the accuracy of the ranking (Table 1).

The ligand core restraint may not be appropriate for all centroids (e.g., Fig. 4F). The reference for the restraint is defined for all receptor configurations by RMSD alignment to the cocrystal structure. Receptor configurations which exhibit large structural differences from the cocrystal structure may have poor binding site alignment which introduces uncertainty into the approach. For some receptor configurations, restraints lead to atypical binding poses with high solvent accessibility.

To test whether using a more complex scoring function and search algorithm at the cost of computational efficiency can improve the predicted rank ordering, we compared between GlideScore SP and GlideScore Extra Precision (XP), with and without the restraint, using the same ensembles from apo MD. Compared to Glide SP, Glide XP (i) has more exhaustive docking by performing Glide SP docking then performing a separate anchor-and-grow sampling procedure, and (ii) the Glide XP scoring function penalizes ligand poses more harshly with desolvation penalties, identification of enhanced binding motifs, and higher receptor-ligand shape complementarity (76). Glide XP has been found to outperform other methods and achieve better drug discovery results than Glide SP (81). We found that Glide XP did not improve our predictions (1). Although the poses predicted by XP were more similar to the cocrystallized poses (4E).

To test whether conformational selection may lead to improved results, we docked to centroids picked from a holo MD simulation. McGovern and Shoichet have showed that use of a holo structure can improve enrichment of lead compound identification (85). We also expected that structures with a ligand would lead to lower ligand core RMSD’s with more accurate active residue positioning. However, the Kendall’s *τ*s of the rank ordering stayed within the same range as the original apo docking, even when the ligand was restrained (1). Upon further analysis of the structural fluctuations of the apo and holo MD centroids, we find that residue PHE71 is restricted by the ligand while other regions of the binding pocket exhibited similar structural variability (Fig. S2). When ligands were blindly docked using Glide SP to the holo structures, the resulting poses remained different than the cocrystal pose. The average RMSD of the docked ligand cores was 5.47 Å from the core of the cocrystal ligand (4C). Overall, the blind docking to structure from the holo MD trajectory had a slightly lower ligand core RMSD compared to the results form docking to the apo MD (Fig. S3). When a core restraint was applied upon docking to configurations from the holo trajectory, even with the influence of the bound ligand on the binding site, the rank ordering did not improve (1).

Although others have suggested that improved poses could yield better scores (86), we found that improvements to the predicted poses from the application of ligand restraints and/or docking to holo receptor conformations did not improve our predictions. This suggests that there may be other confounding factors influencing our results.

## 5 CONCLUSION

In this work we describe our submission to subchallenge 2 of the Drug Design Data Resource (D3R) Grand Challenge 4 where we performed ensemble docking to rank order ligands by binding affinity. We explore and compare several factors including the choice of clustering algorithm for choosing representative receptor conformations and two docking workflows with and without restraints to improve pose accuracy. The different clustering algorithms produce different structural ensembles which can influence the docking results. Owing to the difficulty of docking to the CatS system, which has been recognized by others (87), we find that more sophisticated approaches can improve rank ordering compared to naive settings produced by FRED and GLIDE using a basic ensemble docking workflow (7). Glide yielded better rank order correlations than FRED although no notable differences between the clustering algorithms was observed. We conclude that confounding factors and complications of the CatS system outweigh the benefits of ensemble docking. We explored if rank-order correlation could be improved with better pose accuracy by performing docking with restraints in addition to docking with receptor conformations extracted from a holo trajectory with ligand removed. We find that both approaches improve the pose similarity of docked ligands to related cocrystallized ligands, but do not improve the rank order correlation.

This project illustrates the benefits of partnering with high school and undergraduate students to participate in community challenges. Grand challenges are excellent resources for teaching research skills through a semi-guided, goal-oriented project, with expert curated datasets and deadlines. The students were exposed to important research skills, such as managing time, selecting and performing data analyses, and making publication-quality figures, at early stages of their scientific career. Owing to the computational nature of this challenge, the students also gained experience with data management, computational thinking, and script development. We suggest that student participation in community challenges can benefit both the community and the students and hope this work encourages others to explore this approach.

## AUTHOR CONTRIBUTIONS

Conceptualization, B.C.T., B.R.J., C.T.L. and R.E.A.; Software, J.L.G., D.K.,and C.C.; Investigation, J.L.G., D.K.,and C.C.; Resources, B.C.T., B.R.J., C.T.L. and R.E.A.; Writing – Original Draft, J.L.G, and D.K.; Writing – Review and Editing, J.L.G, D.K., C.C., B.C.T., B.R.J., C.T.L. and R.E.A.; Visualization, J.L.G., D.K.; Supervision and Project Administration, B.C.T., B.R.J., C.T.L. and R.E.A.; Funding Acquisition, R.E.A.

## ACKNOWLEDGMENTS

This work is supported by the National Biomedical Computation Resource NIH Grant P41-GM103426, and the National Science Foundation through The Extreme Science and Engineering Discovery Environment (XSEDE) supercomputing resources provided via Award TG-CHE060073 to R.E.A. C.T.L. is funded by a Hartwell Foundation Postdoctoral Fellowship. We thank D3R and the organizers of Grand Challenge 4 for hosting the challenge and reporting results. We would also like to acknowledge Mason V. Holst, Gaurie Gunasekaran, Gray Thoron, and Jeffery R. Wagner for their contributions to preliminary work and/or helpful discussions.

## SUPPLEMENTARY MATERIAL

The structural similarity of the dataset ligands to cocrystallized ligands, RMSF across receptor structures for the apo and holo trajectories, and comparison of ligand core RMSD across clustering methods are available in the supplemental information.

