## Supplemental Figures for "Benchmarking ensemble docking methods as a scientific outreach project"

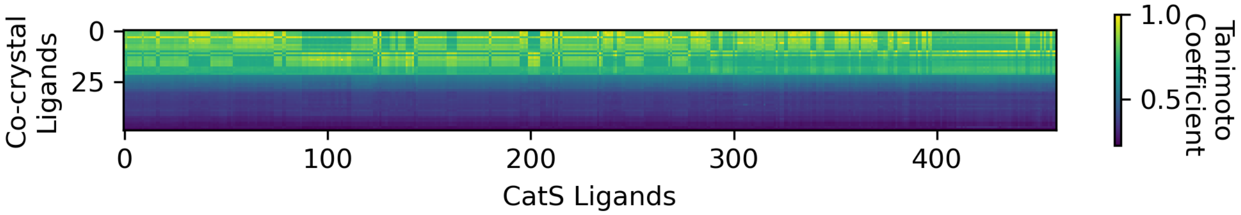


Figure S1: The pairwise Tanimoto coefficients of the cocrystal ligands to the CatS ligands highlights the similarity between the ligands. 48 cocrystal ligands were downloaded from RCSB PDB, and Tanimoto coefficients were calculated using the RDKit Fingerprint implemented in RDKit (1), where a value of 1 represents identical structures and a value of 0 is comparing completely dissimilar structures. The corresponding PDBIDs are in Table S1, and the ligands are ordered by the $L^2$ norm of the Tanimotos, and plotted in matplotlib (2).

Table S1: Corresponding PDB IDs from the RCSB PDB database to cocrystal numbers from Fig. S1. Ordered by the $L^2$ norm of pairwise Tanimoto coefficients between cocrystal ligands and 459 CatS ligands.

| cocrystal Number | PDBID | Ligand ID | $L^2$ Norm |
|---|---|---|---|
| 0 | 5qc8 | BFV | 18.05 |
| 1 | 5qcf | BJD | 17.95 |
| 2 | 5qca | BGJ | 17.9 |
| 3 | 5qc6 | BCJ | 17.79 |
| 4 | 5qci | BJV | 17.64 |
| 5 | 5qcg | BJJ | 17.46 |
| 6 | 5qcd | BHJ | 17.33 |
| 7 | 5qc1 | B9S | 17.31 |
| 8 | 5qc5 | BAJ | 17.13 |
| 9 | 5qcj | BJY | 16.88 |
| 10 | 5qch | BJS | 16.73 |
| 11 | 5qc3 | B9Y | 16.67 |
| 12 | 3iej | 599 | 16.64 |
| 13 | 5qc4 | BC7 | 16.53 |
| 14 | 5qbx | B8V | 16.47 |
| 15 | 5qc0 | BQJ | 16.24 |
| 16 | 5qc2 | BQP | 16.22 |
| 17 | 5qc7 | BQS | 16.22 |
| 18 | 5qcb | BHV | 15.83 |
| 19 | 5qc9 | BG7 | 15.78 |
| 20 | 5qcc | BGY | 15.57 |
| 21 | 5qbu | B8J | 15.28 |
| 22 | 5qbv | N2D | 12.21 |
| 23 | 5qbw | B8S | 11.74 |
| 24 | 5qbz | B8Y | 11.42 |
| 25 | 5qby | N2A | 10.92 |
| 26 | 4p6g | 2FZ | 10.39 |
| 27 | 3n4c | EF3 | 10.23 |
| 28 | 2fye | BCQ | 9.83 |
| 29 | 2h7j | H7J | 9.21 |
| 30 | 2r9o | C28 | 8.35 |
| 31 | 2g7y | MO9 | 8.1 |
| 32 | 2fq9 | CRJ | 8.09 |
| 33 | 2fra | CRV | 8.06 |
| 34 | 2hh5 | GNQ | 7.86 |
| 35 | 2f1g | GNF | 7.76 |
| 36 | 4p6e | 2FC | 7.75 |
| 37 | 5qbz | 935 | 7.72 |
| 38 | 2r9n | Y14 | 7.69 |
| 39 | 2frq | C71 | 7.68 |
| 40 | 2g6d | MQQ | 7.36 |
| 41 | 1npz | C1P | 7.35 |
| 42 | 2fud | CRL | 6.78 |
| 43 | 1nqc | C4P | 6.66 |
| 44 | 2r9o | Y15 | 6.57 |
| 45 | 2r9m | Y11 | 5.97 |
| 46 | 1ms6 | BLN | 5.91 |
| 47 | 5qc1 | O64 | 5.85 |
| 48 | 2op3 | TF5 | 5.1 |

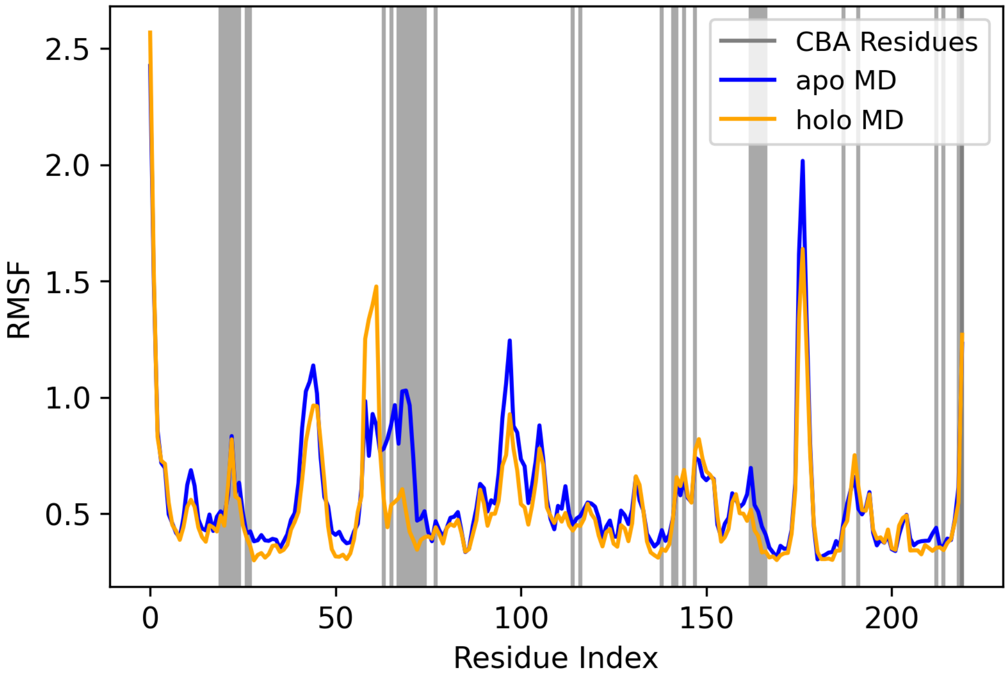


Figure S2: The RMSF of the apo MD and holo MD centroids show slight structural variation in certain residues. RMSF was calculated in CPPTRAJ and plotted using matplotlib (2, 3). The residues that contain any clustering-by-binding-atoms (CBA), determined to be part of the binding site (Fig. 2), are indicated by vertical grey lines, and their indices were extracted with MDTraj (4).

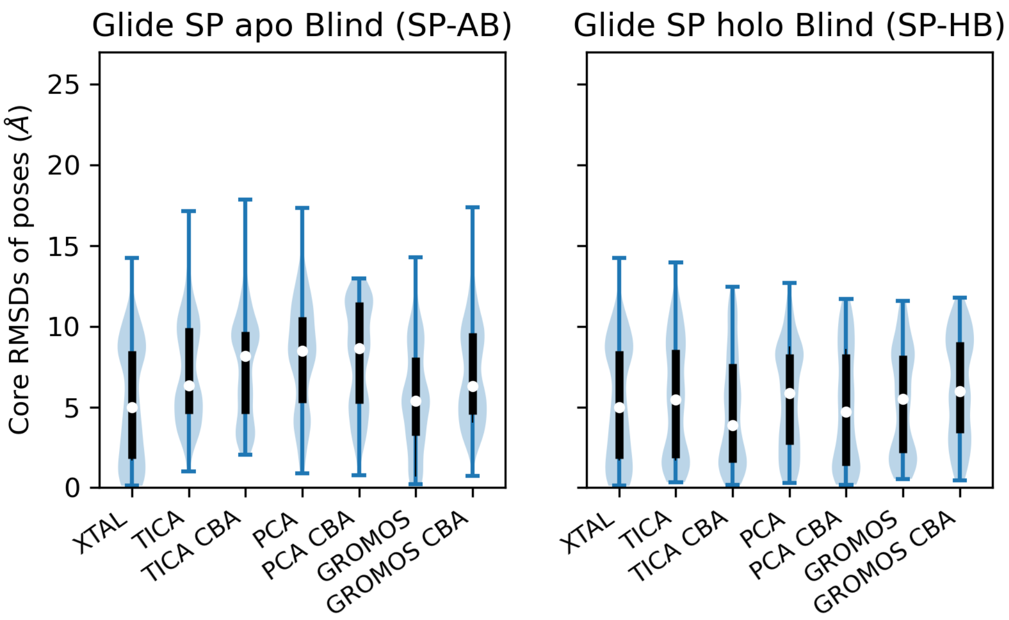


Figure S3: Docking to structures which had previously been docked with a ligand seems to slightly decrease ligand core RMSD. The ligand core RMSDs for the Glide Standard Precision (SP) docking results of the apo blind (SP-AB) and holo blind (SP-HB) are expanded from Fig. 4, into the various clustering methods, where the median seems to be slightly lower when given the holo MD centroids. The ligand core RMSDs were calculated with Schrodinger's python API and plotted in matplotlib (2).
